# Automated identification of aneuploid cells within the inner cell mass of an embryo using a numerical extraction of morphological signatures

**DOI:** 10.1101/2022.09.06.506861

**Authors:** Abbas Habibalahi, Jared M. Campbell, Tiffany C.Y. Tan, Saabah B. Mahbub, Ryan D. Rose, Sanam Mustafa, Kylie R. Dunning, Ewa M. Goldys

## Abstract

**STUDY QUESTION:** Can artificial intelligence distinguish between euploid and aneuploid cells within the inner cell mass of mouse embryos using brightfield images?

**SUMMARY ANSWER:** A deep morphological signature (DMS) generated by deep learning followed by swarm intelligence and discriminative analysis can identify the ploidy state of inner cell mass (ICM) in the mouse blastocyst-stage embryo.

**WHAT IS KNOWN ALREADY:** The presence of aneuploidy – a deviation from the expected number of chromosomes – is predicted to cause early pregnancy loss or congenital disorders. To date, available techniques to detect embryo aneuploidy in IVF clinics involve an invasive biopsy of trophectoderm cells or a non-invasive analysis of cell-free DNA from spent media. These approaches, however, are not specific to the ICM and will consequently not always give an accurate indication of the presence of aneuploid cells with known ploidy therein.

**STUDY DESIGN, SIZE, DURATION:** The effect of aneuploidy on the morphology of ICMs from mouse embryos was studied using images taken using a standard brightfield microscope. Aneuploidy was induced using the spindle assembly checkpoint inhibitor, reversine (n = 13 euploid and n = 9 aneuploid). The morphology of primary human fibroblast cells with known ploidy was also assessed.

**PARTICIPANTS/MATERIALS, SETTING, METHODS:** Two models were applied to investigate whether the morphological details captured by brightfield microscopy could be used to identify aneuploidy. First, primary human fibroblasts with known karyotypes (two euploid and trisomy: 21, 18, 13, 15, 22, XXX and XXY) were imaged. An advanced methodology of deep learning followed by swarm intelligence and discriminative analysis was used to train a deep morphological signature (DMS). Testing of the DMS demonstrated that there are common cellular features across different forms of aneuploidy detectable by this approach. Second, the same approach was applied to ICM images from control and reversine treated embryos. Karyotype of ICMs was confirmed by mechanical dissection and whole genome sequencing.

**MAIN RESULTS AND THE ROLE OF CHANCE:** The DMS for discriminating euploid and aneuploid fibroblasts had an area under the receiver operator characteristic curve (AUC-ROC) of 0.89. The presence of aneuploidy also had a strong impact on ICM morphology (AUC-ROC = 0.98). Aneuploid fibroblasts treated with reversine and projected onto the DMS space mapped with untreated aneuploid fibroblasts, supported that the DMS is sensitive to aneuploidy in the ICMs, and not a non-specific effect of the reversine treatment. Consistent findings in different contexts suggests that the role of chance low.

**LARGE SCALE DATA:** N/A

**LIMITATIONS, REASON FOR CAUTION:** Confirmation of this approach in humans is necessary for translation.

**WIDER IMPLICATIONS OF THE FINDINGS:** The application of deep learning followed by swarm intelligence and discriminative analysis for the development of a DMS to detect euploidy and aneuploidy in the ICM has high potential for clinical implementation as the only equipment it requires is a brightfield microscope, which are already present in any embryology laboratory. This makes it a low cost, a non-invasive approach compared to other types of pre-implantation genetic testing for aneuploidy. This study gives proof of concept for a novel strategy with the potential to enhance the treatment efficacy and prognosis capability for infertility patients.

**STUDY FUNDING/COMPETING INTEREST(S):** K.R.D. is supported by a Mid-Career Fellowship from the Hospital Research Foundation (C-MCF-58-2019). This study was funded by the Australian Research Council Centre of Excellence for Nanoscale Biophotonics (CE140100003), the National Health and Medical Research Council (APP2003786) and an ARC Discovery Project (DP210102960). The authors declare that there is no conflict of interest.

## 1. INTRODUCTION

Mitotic errors during preimplantation embryo development can result in the emergence of a subset of aneuploid cells, which, at high levels may threaten the achievement of a successful pregnancy with healthy offspring (Bolton, et al., 2016). As such, the transfer of embryos with a major degree of aneuploidy is highly undesirable in assisted reproductive technologies. Preimplantation genetic testing for aneuploidy (PGT-A) has consequently received widespread adoption. However, the invasiveness of this technology, which most commonly involves the collection and karyotyping of a small biopsy of trophectoderm cells from the preimplantation embryo prior to transfer, creates major drawbacks, including a potential increased risk of preeclampsia (Mastenbroek, et al., 2011). Additionally, trophectoderm biopsy has the potential to impact implantation, embryo survival, clinical pregnancy and live-birth rates (Rubino, et al., 2020). In principle, this is offset by the improved outcomes of not transferring aneuploid embryos, however the impact remains, and a non-invasive methodology of karyotyping is highly desirable. Furthermore, the karyotype of trophectoderm cells – which contribute to the placenta – will not always be a direct reflection of the composition of the inner cell mass (ICM) cells – specifically the epiblast – which gives rise to the fetus. As such, techniques based on trophectoderm biopsy have the potential to give rise to both false positive and negative findings (Vera-Rodriguez and Rubio, 2017).

Genetic assessment of cell-free DNA (cfDNA) delivered by embryos into the culture media is a proposed technology for non-invasive preimplantation genetic testing of aneuploidy (niPGT-A) (Rubio, et al., 2019). Although elevated concordance with assessments based on trophectoderm biopsy has been reported in some instances (Feichtinger, et al., 2017, Huang, et al., 2019), other authors have found variability (Vera-Rodriguez, et al., 2018, Xu, et al., 2016). This is hypothesized to arise from embryos having a varying degree of aneuploidy (Xu, Fang, Chen, Chen, Xiao, Yang, Wang, Song, Ma, Bo, Shi, Ren, Huang, Cai, Yao, Xie and Lu, 2016) and contamination of medium by maternal DNA (Vera-Rodriguez, Diez-Juan, Jimenez-Almazan, Martinez, Navarro, Peinado, Mercader, Meseguer, Blesa, Moreno, Valbuena, Rubio and Simon, 2018). As such, there is a risk that any diagnosis – positive or negative – will not genuinely reflect the ploidy status of the ICM (Gleicher and Barad, 2019).

To address the need for the direct assessment of ICM ploidy, we applied artificial intelligence to the assessment of brightfield images of the ICM of blastocysts. Recently, AI has been extensively employed to characterize biomedical and clinical images for monitoring and diagnosing disease. This allows the precise personalised treatment schedules with excellent accuracy and sensitivity and also to identify detailed imaging changes due to biological alterations (Louis, et al., 2021, Pesapane, et al., 2018). Attempts have been made to employ AI for embryo assessment, however results have been variable (Bartolacci, et al., 2022, Patil, et al., 2018, VerMilyea, et al., 2020) and analysis has mostly relied on expensive time lapse imaging (Bormann, et al., 2020, Javadi and Mirroshandel, 2019, Kandel, et al., 2020, Khosravi, et al., 2019, Liao, et al., 2021, Tran, et al., 2019).

Here, we identified a deep morphological signature (DMS) of embryo images, comprising multiple image features. Using blastocysts with aneuploidy induced by the spindle assembly checkpoint inhibitor, reversine, this paper shows that our DMS is capable of detecting morphological changes caused by aneuploidy. In order to produce this DMS, brightfield images of cells or ICM with aneuploidy induced by reversine treatment and confirmed by sequencing were isolated and morphological features of the embryo ICM such as shapes and textures were obtained by three specially designed deep learning nets. These had a unique construction designed to extract precise data-driven information from a limited number of images. Deep convolutional learning by these nets was then followed by discriminative analysis, and feature selection by the swarm intelligence approach. To ensure that our technique is generalisable, cross-validation was employed during DMS discovery, which also minimizes the potential for overfitting. Our advanced artificial intelligence method improves on traditional image feature analysis and makes it possible to non-invasively identify ICM ploidy in early embryos, employing only equipment used in routine embryo inspection.

## 2. MATERIALS AND METHODS

### 2.1. Samples and preparation

Cell culture, animal procedures and sample preparation were carried out as described in Reference (Tan, et al., 2022). University of Adelaide’s Animal Ethics Committee approved the experiments, and they were carried out in accordance with the Australian Code of Practice for the Care and Use of Animals for Scientific Purposes. In brief, three-week-old (female) and 6–8-week-old (male) CBA x C57BL/6 F1 from Laboratory Animal Services (LAS; University of Adelaide, SA, Australia) were kept in in a 12h light/dark cycle with *ad libitum* chow and water. Follicle development was stimulated by 5 IU equine chorionic gonadotrophin (eCG; Braeside, VIC, Australia) followed 48 h later with 5 IU (i.p.) human chorionic gonadotrophin (hCG; Kilsyth, VIC, Australia) to stimulate ovulation, and then culling and cumulus oocyte complex (COC) collection 14 h after that. Male mice were culled one hour prior to IVF, spermatozoa were released by blunt dissection and allowed to capacitate for 1 h. Mature COCs were co-cultured capacitated spermatozoa for 4 h and the now presumptive zygotes were cultured in groups of 10 for 24 h with cleaved embryos being left for further development. All gamete and embryo culture was carried out in media at 37°C, under paraffin oil, in a humidified atmosphere (5% O_2_, 6% CO_2_, balance of N_2_).

Embryo aneuploidy was generated by treating mouse embryos with reversine, a reversible spindle assembly checkpoint inhibitor, (reversine 0.5 μM) from the 4- to 8-cell stage. At the blastocyst-stage the ploidy status of the ICM was confirmed by dissection of the ICM followed by sequencing (as described in Tan et al 2022). Only blastocysts whose ICMs were confirmed to contain some degree of aneuploid cells were included in subsequent analyses. Imaging of blastocysts was carried out in glass-bottomed dishes (Ibidi, Martinsried, Planegg, Germany) containing pre-warmed Research Wash Medium under paraffin oil. For the analysis of aneuploid cells, primary human fibroblasts (Coriell Institute for Medical Research, USA) with known karyotypes (46, XY (GMO0970); 46, XX (GMO3525); triploid (GM01672); trisomy: 13 (GM02948), 18 (GM01359), 21 (AG06922), XXX (GM04626), and XXY (GM03184)) were plated and cultured overnight at 1 x 10^5^ in 35 mm glass bottom dishes (Ibidi, Martinsried, Planegg, Germany). To determine if reversine had an effect on cell morphology not specific to aneuploidy, we treated aneuploid human fibroblast cells (GM01359 and AG06922) with reversine (0.5 μM for 8 h).Following this, cells were washed three times and allowed to recover for 2.5 h in culture media. Prior to imaging, cells were washed twice in PBS without calcium and magnesium and imaged in Hank’s balanced salt solution (Gibco, ThermoFisher Scientific, Waltham, MA, USA).

### 2.2. Bright field imaging

Bright field (BF) microscopy was performed using an Olympus iX83 microscope with a 60× oil (NA 1.15) objective. Images were captured by an electron multiplying CCD (NuvuTM 1024, Canada) with 1024×1024 pixel (13×13 mm). Bright field imaging used here employed exposure times of a fraction of second. Cells and ICMs of embryos – which had been specifically focused on during imaging – were manually segmented prior to analysis.

### 2.3. Data analysis

Our data analysis extracted comprehensive morphological information from BF images taken of cells and the manually segmented ICM regions of blastocysts. The data analysis schematics (Figure 1) illustrates our analysis approach. We used external and internal cross validation for our analysis. The external cross validation was used for the DMS discovery, and the internal cross validation was used for classifier training. First, to facilitate external cross validation, we divided data into training (75%) and testing (25%) data sets(Habibalahi and Safizadeh, 2014). We then applied image augmentation (see section 2.3.1) and a deep convolutional learning approach (LeCun, et al., 2015) where several (N=3) architecturally different deep learning nets(Habibalahi, et al., 2021, Habibalahi, et al., 2022) each of which with specific resolutions were employed to generate precise and data-driven image information (details of the nets are given in section 2.3.2). The DMS was discovered through iterative use of swarm intelligence (Kennedy, 2006) which utilizes cooperative behavior of a number of self-organizing, decentralized, naïve agents to efficiently achieve optimum results (Blum and Li, 2008, Garnier, et al., 2007). In this part (see section 2.3.3) we employed a Fisher distance gauge function to assess feature subsets quality offered by swarm intelligence in an iterative manner (Beni, 2009, Habibalahi, et al., 2022, Hu, et al., 2004, Panigrahi, et al., 2011) that eventually converge to select an optimised set of features. The final DMS employed to train a classifier which is support vector machine (SVM) as our preferred classifier (Furey, et al., 2000) in this study. SVM is a powerful supervised technique capable of dealing with biological data with sparse condition and. SVM has limited overfitting risk since it uses linear predictor function to distinguish the label of data(Habibalahi, et al., 2022). The SVM was cross validated internally using standard k-fold cross validation (k=10) and the classification performance was assessed by receiver operating characteristic (ROC) curve and associated area under the curve (AUC)(Habibalahi, 2019, Habibalahi, et al., 2019).

**Figure 1.**
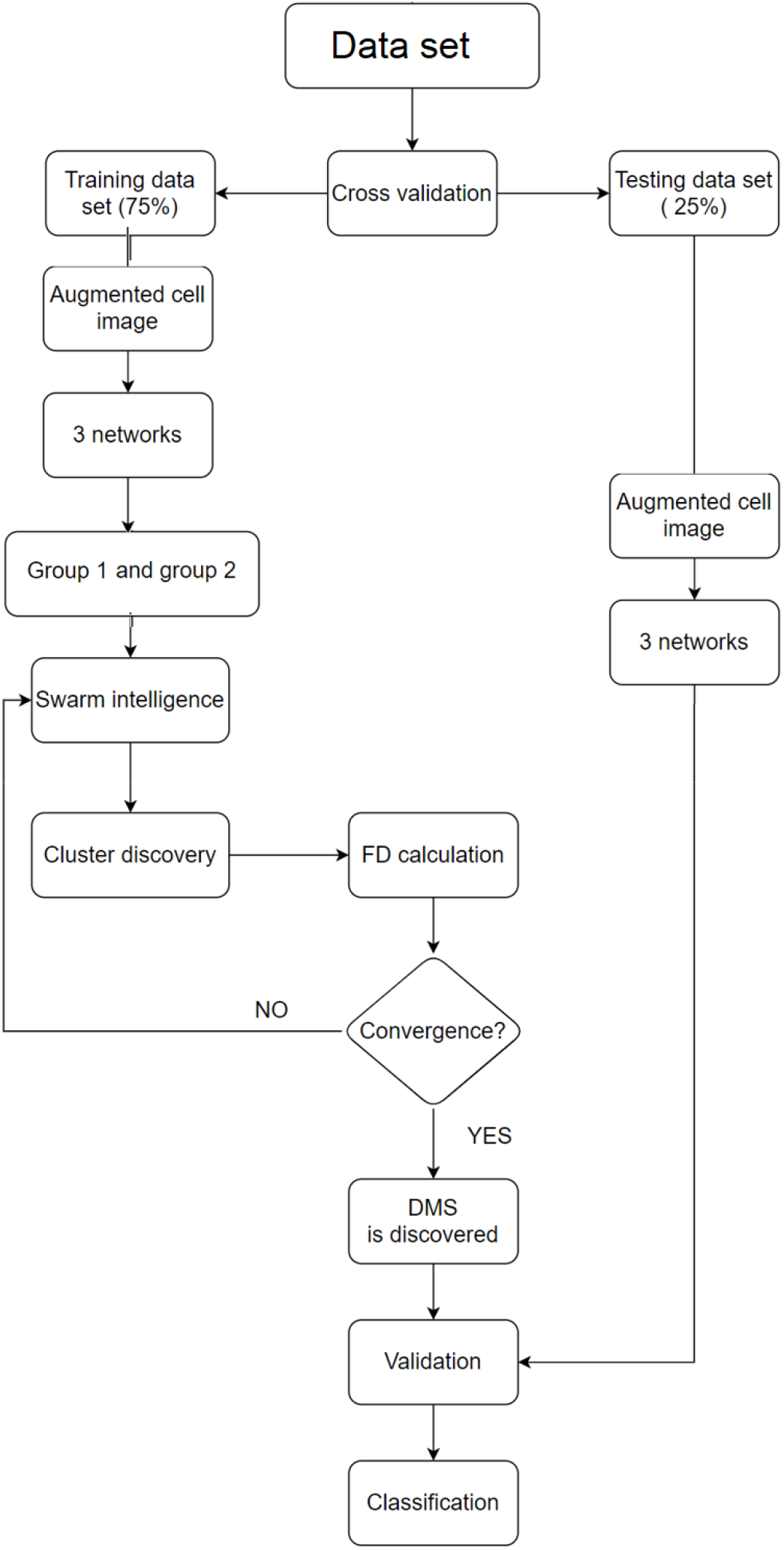
Data analysis methodology (details are described in section 2.3).

#### 2.3.1. Image augmentation

Image augmentation is a standard method in AI (Wang and Perez, 2017) to enrich the data set artificially through inserting images intuitively comparable to the original images (here embryos images). Original images rotated by several angles, or mirrored along different axes – since the positioning of embryo images on the microscope is not appropriate to karyotype(Wang and Perez, 2017). Image augmentation also expands the data set which improves the deep learning performance by enhancing training images to discover stronger DMS. Embryo images and their corresponding reflections, in this study, were rotated at various angles (45°, 90°, 135°, 180°) (Habibalahi, Bertoldo, Mahbub, Campbell, Goss, Ledger, Gilchrist, Wu and Goldys, 2021, Habibalahi, Campbell, Mahbub, Anwer, Nguyen, Gill, Wong, Pollock, Saad and Goldys, 2022).

#### 2.3.2. Deep convolutional neural networks

A deep convolutional neural network (CNN) approach was employed in our study for feature discovery. To identify features, the CNN uses convolutional layers successively applied to an input image while in traditional algorithms features were obtained by application of mathematical definition of features (Campbell, et al., 2019). A sequence of three nets (Net1, Net2 and Net3) were designed for this work(Habibalahi, Bertoldo, Mahbub, Campbell, Goss, Ledger, Gilchrist, Wu and Goldys, 2021, Habibalahi, Campbell, Mahbub, Anwer, Nguyen, Gill, Wong, Pollock, Saad and Goldys, 2022). Each CNN generated a separate feature collection and subsequently all features were collated. Each net has specific configurations and resolutions, which leads to detailed and comprehensive image feature extraction(Habibalahi, Bertoldo, Mahbub, Campbell, Goss, Ledger, Gilchrist, Wu and Goldys, 2021, Habibalahi, Campbell, Mahbub, Anwer, Nguyen, Gill, Wong, Pollock, Saad and Goldys, 2022). Net 1 has a deep depth including 153 convolutional layers whose filters were taken from ResNet (He, et al., 2016). Net 2 has medium depth containing 22 convolutional layers separated from GoogLe net (Szegedy, et al., 2015) and Net 3 has shallow depth only with 6 convolutional layers secured from the Krizhevsky net (Krizhevsky, et al., 2012). To finetune each net for our specific study, the last layer of each CNN was retrained by the data set from this study. Employing these three nets, ~7000 deep features were extracted for each cell or embryo, producing this study dataset.

#### 2.3.3. Swarm intelligence and discriminative analysis

Swarm intelligence is an artificial intelligent approach stimulated by the collective behaviour immature information-processing cooperating agents (Kennedy, 2006). This approach was detailed in our recently published paper(Habibalahi, et al., 2020). Briefly, the simple artificial agents (here feature subsets) are moving satisfying a pre-defined development rule by maximizing Fisher Distance (FD) as a measure of optimization. Initially, an agent population (here feature subsets) are created, and corresponding FD is calculated for each agent(Habibalahi, Campbell, Mahbub, Anwer, Nguyen, Gill, Wong, Pollock, Saad and Goldys, 2022). Next, agents are continually revised based on the pre-defined strategy of evolution till satisfying the convergence of FD. In this study, swarm intelligence used for selecting features while a number of agents (N=50) were feature subset candidate of entire features produced by three CNNs (Nf~7000) (Kennedy, 2006). The principle in the swarm intelligence procedure was to maximize the FD (Tsuda, et al., 2003, Vapnik, 2013). To assess FD, the data from groups under consideration (euploid and aneuploid cases) were reflected on a 2-D discriminative space. This discriminative space was generated based on two canonical variables maximizing group separation.

## 3. RESULTS

### 3.1. DMS detects a significant morphological difference in cells associated with aneuploidy

To investigate the plausibility of identifying aneuploidy using DMS, we initially used primary human fibroblast cells with confirmed karyotypes for aneuploidies (trisomy: 21, 18, 13, 15, 22, XXX and XXY) compared to primary, euploid cells from two sources. A cluster separation plot was created (Fig 2a) to illustrate the separation of euploid-aneuploid classes in the augmented dataset. The observed separation was validated using a testing dataset (Figure 2 (b)) where closely similar degree of separation was achieved. Further, the DMS was fed to a SVM classifier which achieved an AUC= 0.89 and accuracy = 82%, for the separation of euploid and aneuploid cells (Figure 2 (c)). This demonstrates that there are cellular features detectable by DMS, and common across multiple forms of aneuploidy, which can be used to classify aneuploid and euploid cells. To investigate the impact of reversine on morphology (a potential confounder of the ICM DMS described below), aneuploid fibroblasts were treated with reversine. In Figure 2 d their featured were projected onto the DMS space. We found that the DMS continued to sort them with the endogenously aneuploid cells, demonstrating that the reversine treatment is not having a morphological impact beyond the induction of aneuploidy. and supports reversine’s impact on cell morphology being the result of the induction of aneuploidy as opposed to other potential molecular changes.

**Figure 2.**
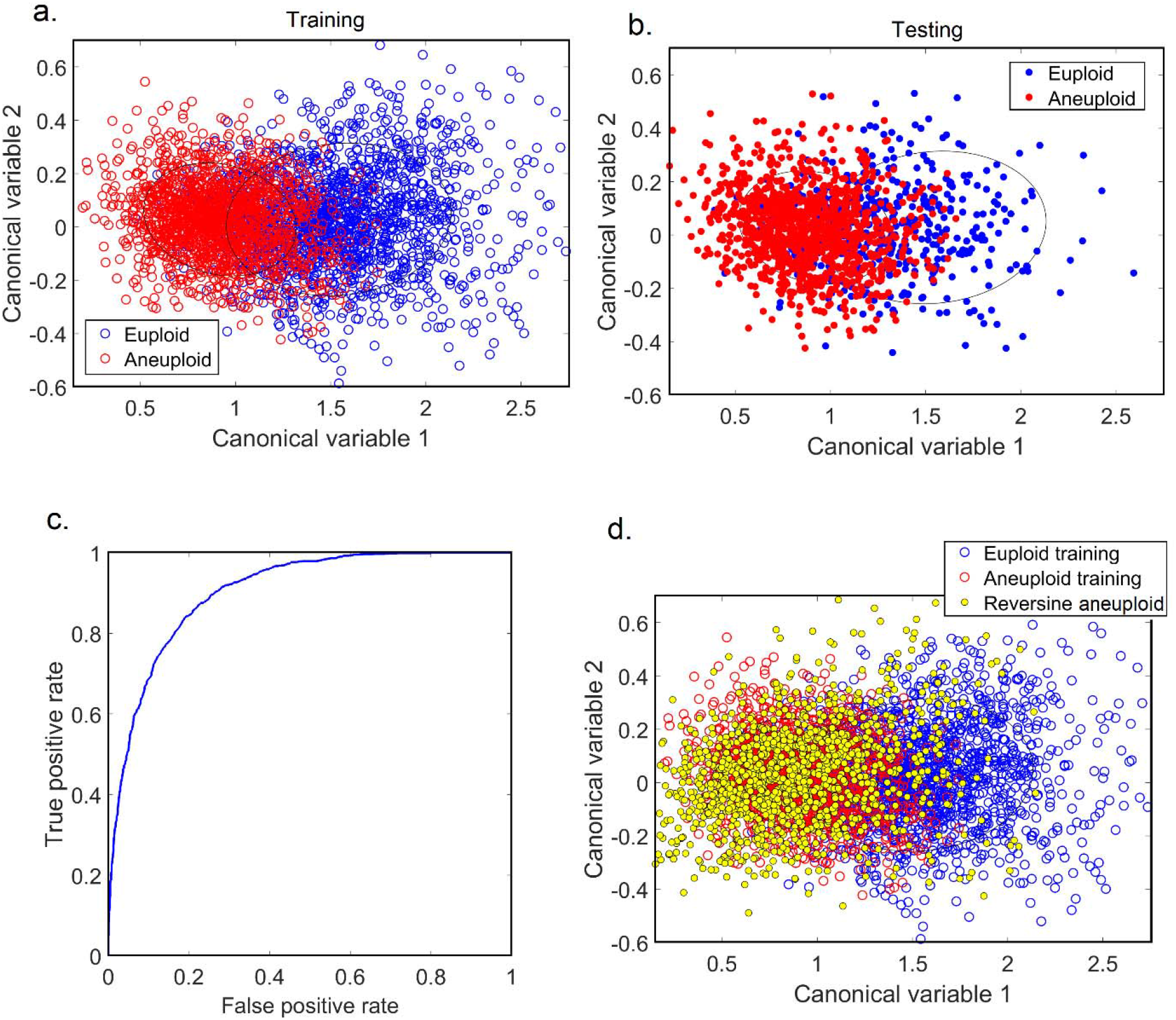
DMS for aneuploid and euploid fibroblast cells. A DMS was developed per the analytic approach defined in materials and methods. A graph of cluster separation was created where each data point shows the final product of image augmentation (a, blue: male and female euploid cells; red: Triploid and Trisomies 13, 18, 21, XXX, XXY). n = 2184 for augmented euploid cells; n= 2400 for augmented aneuploid cells. Canonical values are specific linear combinations of DMS features. (b). Testing data points were projected on the discriminative space created by training data points. (c) Receiver operating characteristic (ROC) curve was produced in order to establish the separation accuracy of aneuploid and euploid cells (AUC = 0.89). (d) reversine-treated aneuploid fibroblasts (yellow, n=1200) projected into the DMS space with aneuploid and euploid fibroblast cells (same as (a).

### 3.2. Aneuploid and euploid ICMs display different morphological properties

We next assessed the morphological properties of ICMs with confirmed aneuploidy. We found that the presence of aneuploidy had a consistently strong impact on blastocyst ICM morphology. This is demonstrated in Figure 3a, where two distinct clusters were formed (as apparent in the absence of intersection of ellipses drawn one standard deviation around the mean value) which was validated with testing data points (Figure 3b). The SVM classifier ROC is shown in Figure 3 b whose performance is very high, with AUC=0.98+/− 0.01. This indicates high sensitivity to the presence of aneuploidy within the ICM. Further investigation showed that the ICM size was significantly reduced by aneuploidy condition (Figure 3 d) with Feature 7 of the DMS correlating closely with ICM size (R^2^=0.73).

**Figure 3.**
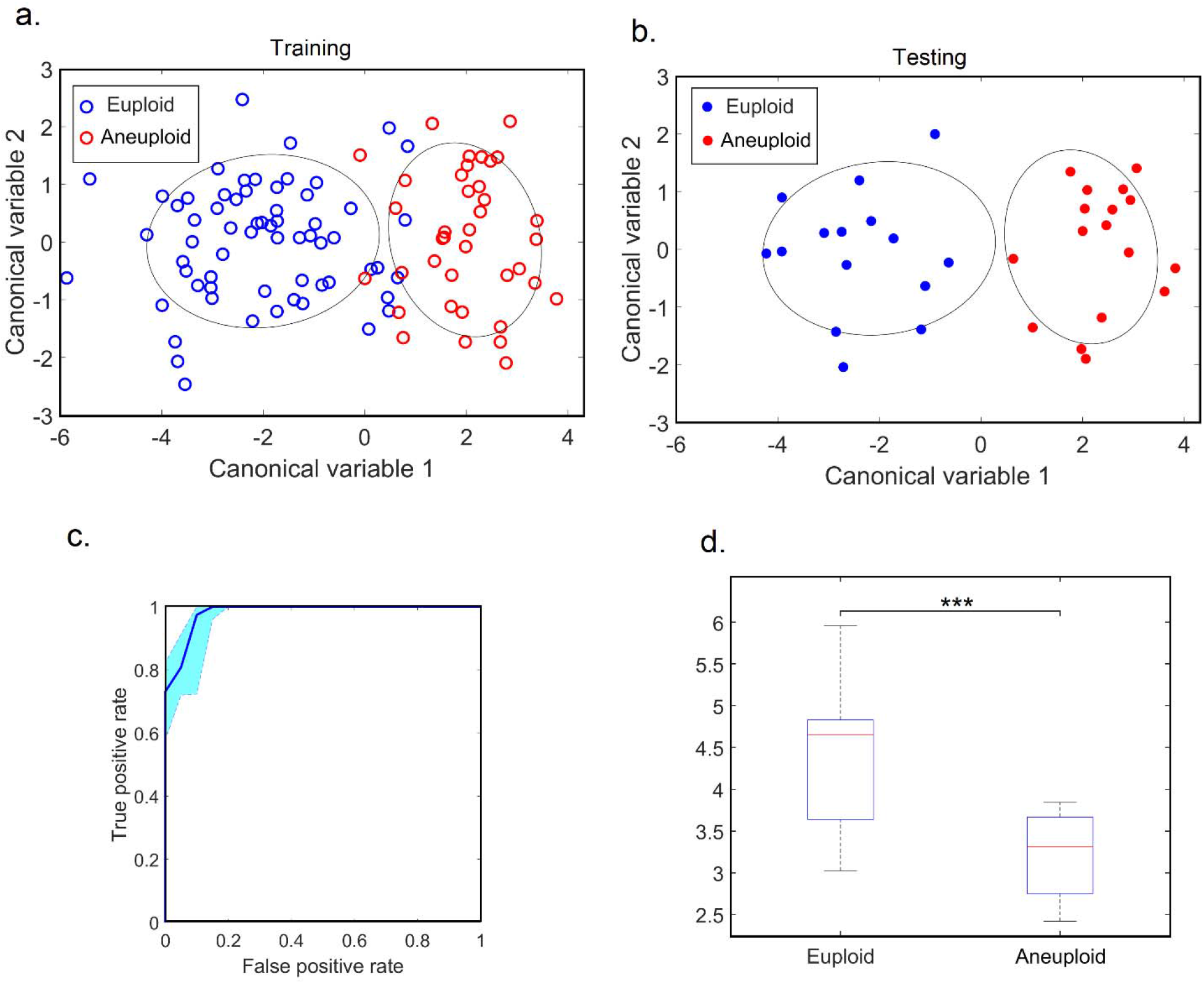
DMS for euploid and aneuploid ICMs (n=78 for augmented euploid ICM; n= 54 for augmented aneuploid ICM from 13 and 9 images, respectively). (a) euploid and aneuploid cluster separation (training data set, 75% of data) (b) euploid and aneuploid cluster validation on testing data points (25% of data) (c) SVM Classifier ROC curve to define embryo labels (due to comparatively small number of embryo images, sampling error was obtained using standard bootstrapping method (Efron and Tibshirani, 1994) with 100 resampled number to obtain 90% confidence margin as shown with light blue color) (d) Size of ICM univariate feature analysis (*** represents p<0.01).

## 4. DISCUSSION

Embryo quality is directly related to pregnancy success rate (Zhu, et al., 2014), and therefore noninvasive and cost-effective procedures for assessing its quality are needed. Currently, the standard, noninvasive methodology for selecting quality embryos is by the inspection visible morphological and developmental features, such as cytoplasm granularity, zona thickness, oolemma roundness, cytoplasm color, blastomere number, multinucleation, degree of fragmentation, blastocoel size, timing of cleavage and morphokinetics, and more (Curchoe and Bormann, 2019). Correlations between these embryo morphology features, as scored by embryologists, and the presence of aneuploidy have been demonstrated (Majumdar, et al., 2017, Minasi, et al., 2016), however they are not strong enough for general clinical implementation. This is likely a result of subjective manual grading and also only considering aspects of morphology that are perceivable and recognizable through human perception, which uses only a small subset of potentially available diagnostically relevant features.

To address this need, we have applied automated morphological analysis based on AI to brightfield images focused on the ICM of blastocyst-stage embryo. Brightfield imaging is a basic microscopy technique routinely used in the clinic to inspect oocytes, embryos and other reproduction materials (REF). To standardise image classification, a DMS was developed in this study using a combination of AI methods including swarm intelligence, deep learning and discriminative analysis, to differentiate aneuploid from euploid ICMs based on brightfield images. DMS can be extracted automatically, and it recognizes image variations resulting from features detectable by human inspection, including ICM size. Besides, this method is able to identify features imperceptible through conventional visual examination of embryos, such as the detailed spatial pixel distribution within an image as well as pixel interrelationships (Guo, et al., 2016).

In this study, DMS application was initially demonstrated by a successful discrimination of euploid fibroblasts from karyotyped aneuploid fibroblasts with a high classification performance (AUC= 0.89). Further, aneuploid fibroblasts were treated with reversine, but no changes were induced that caused them to be were not distinguishable from endogenously euploid lines (Figure 2d). As well as reinforcing the sensitivity of the DMS to aneuploidy, this result supports the validity of the latter DMS developed for detecting aneuploidy in ICMs. An AUC of 0.98 represents very strong discrimination of aneuploid from euploid ICMs. The size of the ICM was also significantly impacted by aneuploidy (Fig. 3d). In addition, our DMS achieved higher accuracy than comparable published models on blastocysts (Barnes, et al., 2020, Huang, et al., 2021, Miyagi, et al., 2019, Ortiz, et al., 2021). Other published studies have indicated that clinometric data can further strengthen prediction which should be incorporated into future clinical applications (Barnes, Malmsten, Zhan, Hajirasouliha, Elemento, Sierra, Zaninovic and Rosenwaks, 2020, Ortiz, Morales, Lledo, Garcia-Hernandez, Cascales, Vicente, González, Ten, Bernabeu and Llácer, 2021). We recently demonstrated that hyperspectral imaging of cell autofluorescence was able to detect aneuploidy in in vitro cultured mouse blastocysts (Tan, Mahbub, Campbell, Habibalahi, Campugan, Rose, Chow, Mustafa, Goldys and Dunning, 2022). However, this technology requires specialised equipment for the collection of images, while the collection of brightfield images can be performed with standard microscopy equipment found in any embryology lab. As such there is a significant translational advantage to this approach.

Although the DMS which we have developed for mouse ICMs cannot be applied directly to human blastocysts, the strategy described here has high potential for clinical implementation as it would not require any specialist or expensive equipment beyond standard brightfield microscopy used in routine embryology. As the proof of concept, this study is the first step for DMS development to enhance the treatment efficacy and prognosis capability for infertility patients. Our technology employs simple bright field imaging to develop DMS, which is a low cost and non-invasive approach compared to other types of pre-implantation genetic testing to detect aneuploidy. Future work will include investigating its application to human embryos in order to build a DMS able to be applied clinically to maximise the likelihood of successful pregnancy and healthy offspring.

## AUTHORS’ ROLES

A.H, E.G, K.R.D. conceived the idea for the study. T.C.Y.T., J.M.C., E.M.G. and K.R.D. were involved in experimental design. A.H, T.C.Y.T., J.M.C., S.M., E.M.G. and K.R.D. were involved in interpretation of data. T.C.Y.T., S.B.M., J.M.C., A.H.,, R.D.R. and. were involved in data acquisition and data analysis.

A.H. performed artificial intelligence, swarm intelligence, discrimination analyses and biostatistics. A.H and J.C prepared the manuscript. A.H, J.C, E.M.G., T.C.Y.T. and K.R.D. critically edited the final version of the manuscript. E.G and K.R.D supervised the study. All authors critically revised and approved the final version of the manuscript.

## ACKNOWLEDGEMENTS

Personal acknowledgements here if any.

